# Goals Drive Adaptive Analysis of Linguistic Features During Reading

**DOI:** 10.64898/2026.09.25.754337

**Authors:** Pepita Alex, Julia Schwarz, Benjamin W. Tatler, Agnieszka E. Konopka, Anastasia Klimovich-Gray

**Author notes:** **Correspondence:** Anastasia Klimovich-Gray Address: William Guild Building, King’s College, University of Aberdeen, Aberdeen AB24 3FX.

## Abstract

Reading behaviour is sensitive to contextual demands. For example, when reading to study or answer questions, readers strategically allocate more attention to goal-relevant sections, read slower, and regress more frequently to extract specific information based on task demands. This suggests that readers flexibly adapt their strategies according to goals and situational constraints. However, the neurocognitive mechanisms supporting this flexibility remain unclear. To test this, we combined eye-tracking with EEG during naturalistic reading and examined how short-term reading goals modulated the neural tracking of hierarchical linguistic features at the orthographic (orthographic neighbourhood size), lexical (word frequency), and contextual-semantic (Surprisal) levels. Forty native English speakers read narratives in three goal conditions: Baseline (no goal), Proofreading (detect spelling errors), and Comprehension (attend to content). Fixation-aligned word-level EEG activity was modelled using multivariate temporal response function to estimate the strength of feature-specific neural tracking. Orthographic tracking was significantly enhanced during Proofreading relative to Baseline, and predicted better Proofreading accuracy. Conversely, stronger frequency tracking predicted better Comprehension accuracy. Contextual-semantic tracking was not modulated by task goal. These findings suggest that reading goals selectively bias neural processing towards linguistic features (sublexical vs. lexical) that support successful task performance, providing evidence for short-term functional plasticity in linguistic processing and supporting a flexible predictive processing account of goal-directed reading.

**Highlights:**

- Cortical analysis of text is flexibly modulated by individual’s goals
- Proofreading selectively facilitates cortical tracking of orthographic information
- Reading for context does not enhance tracking of contextual-semantic features
- Neural tracking of goal-relevant features predicts better task performance

---

Recent neurocognitive models of language comprehension increasingly view language analysis as dynamic and computationally flexible, where processing of hierarchical linguistic features and representational units can be flexibly up- or down-regulated by attentional and contextual demands (e.g., Dogonasheva et al., 2025; Mattys et al., 2026). On the mechanistic level, the ability to up- or down-regulate the analysis of hierarchical linguistic features can be implemented within the predictive processing framework (Clark, 2013; Friston, 2005; Friston, 2010). According to this framework, bottom-up prediction errors (i.e., mismatch between prediction and input) are propagated to update higher-level internal models when incoming input deviates from expectations. Importantly, the degree of model-updating also depends on dynamic precision weighting, which determines the relative influence of top-down priors versus bottom-up incoming input during inference across the levels of the processing hierarchy (Clark, 2016; de Lange et al., 2018; Feldman & Friston, 2010; Friston, 2018; Friston et al., 2017; Hohwy, 2020). Processing levels that are assigned greater precision exert a stronger influence during inference, whilst those with weaker precision are down-weighted. Current goals can dynamically shape weighting of information across the processing levels to optimise performance to the task. Therefore, precision weighting provides a framework for the integration of linguistic information in a way that is consistent with current contextual and higher-order goals. However, the majority of the proposed models that include such goal-adaptation capacity focus on spoken language. Naturalistic reading is a highly goal-sensitive process (van den Broek et al., 2001; Kaakinen & Hyönä, 2007); yet we know very little about whether and how flexible analysis of linguistic information is achieved on the neural level during text analysis.

Behavioural evidence suggests that text analysis is highly sensitive to contextual demands, with readers flexibly adapting their strategies according to overarching goals and situational constraints (Lorch et al., 1993; Paris et al., 1983; van den Broek et al., 2001).

Longer fixation durations generally reflect greater cognitive effort and/or processing load (Rayner, 1998), and these are systematically modulated by reading goals and task contexts, demonstrating that eye movements are influenced by contextual relevance rather than reflecting fixed oculomotor routines (Kaakinen & Hyönä, 2010). For example, when reading to study, to search for information, or to answer questions, readers strategically allocate more attention to goal-relevant sections of text, read more slowly, and regress more frequently to extract specific information based on current task demands (Kaakinen & Hyönä, 2007; Kaakinen et al., 2003). At a more fine-grained information-processing level, different goals have also been shown to bias the depth and locus of linguistic analysis of text. Readers adopt distinct strategies depending on the level of information they seek to extract: compared to reading for comprehension, proofreading typically leads to slower and more careful reading, with increased sensitivity to word length and frequency effects, potentially reflecting increased orthographic and lexical processing (e.g., Daneman et al., 1995; Kaakinen & Hyönä, 2010; Schotter et al., 2014). Importantly, these effects occur even in the absence of actual errors (Kaakinen & Hyönä, 2010); this indicates that the goal itself modulates level-specific processing rather than a general error detection mechanism (i.e., not driven by bottom-up stimulus properties). In contrast, word detection tasks that do not require comprehension (e.g., scanning/search) selectively reduce lexico-semantic engagement while increasing orthographic processing, resulting in poorer comprehension and diminished/absent word frequency effects (e.g., Rayner & Fischer, 1996; Radach et al., 2008). Together, these behavioural findings indicate that readers can dynamically re-distribute processing resources across processing levels depending on task demands. From a predictive processing perspective, changes in reading strategy can be explained as goal-dependent modulation of precision weightings guided by higher-level metacognitive priors. Crucially, these priors do not merely encode expectations about linguistic content but about which reading strategy is appropriate in a given context (Lorch et al., 1993), for example, on whether to prioritise higher-level semantic versus lower-level orthographic processing.

However, there is little neurocognitive evidence of whether and how goals shape neural computations that guide reading behaviour in real time. The majority of the cited work utilises eye movements as behavioural proxies of cognitive processing. While these measures have provided valuable insight into how reading behaviour adapts to contextual demands, they cannot directly specify the underlying neurocognitive mechanisms. For instance, although fixation and attention are tightly coupled during reading, where attention is typically allocated to the fixated word (Inhoff & Briihl, 1991; Rayner, 1998; Rayner, 2009), fixation alone does not guarantee meaningful semantic extraction (e.g., Reichle et al., 2010). Covert shifts of attention can occur regardless of overt gaze (e.g., Posner, 1980), and orthographic and semantic facilitation can occur parafoveally, with processing continuing after fixation offset (Rayner, 1998; Rayner et al., 1989; Reichle et al., 2003; Schotter et al., 2012; Wang et al., 2025). As a result, identical eye-movement patterns may arise from different cognitive operations, and conversely, similar cognitive states may produce different reading behaviours. Inferring goal-direct adaptation in computations at distinct levels of linguistic analysis solely from eye-tracking data is therefore limited.

## Present study

The present study addresses these gaps through combining EEG with eye-tracking during naturalistic goal-oriented reading. This allows us to move beyond surface-level behavioural indices to directly characterise whether and how task goals modulate the neural tracking of different linguistic features in real time and in conditions that closely approximate everyday reading. In a within-subjects design, participants completed a reading task with three goal conditions: Baseline (no goal instructions), Proofreading (instructed to detect misspelled words), and Comprehension (instructed to attend to story content). Consistent with the precision-weighting account, we hypothesised that task goals should alter the distribution of precision across processing levels, selectively biasing processing towards features most relevant to the current goal while reducing engagement on goal-irrelevant processing to accommodate immediate demands under limited cognitive resources.

To address this, we require analytical approaches capable of modelling the relationship between stimulus features and neural responses during naturalistic reading whilst considering potential confounds (see Dimigen & Ehinger, 2021). To this end, we adopted multivariate temporal response function (mTRF) modelling (Brodbeck, et al., 2023; Crosse et al., 2016; Crosse et al., 2020), which has been widely applied to naturalistic continuous speech paradigms (e.g., Gillis et al., 2021; Theron-Grimaldi et al., 2025), and very recently, to continuous text processing via rapid serial presentation (Soghoyan et al., 2025). mTRF allows simultaneous modelling of multiple predictors and estimates the linear relationship between stimulus features and multi-channel neural activity over time. By applying a sliding temporal window (time-lags), mTRF generates temporal response functions (TRFs) that predict the latency and profile of the brain’s response to specific features in real time. Model performance is quantified using correlation coefficients (R-values), which index the correlations between predicted and observed neural responses. Higher (partial) R-values therefore reflect stronger and more robust neural tracking of a given feature. Systematic differences in tracking strength (R-values) across the three goal conditions may reflect shifts in the relative influence of particular processing levels.

Modelling the neuro-cognitive mechanisms of naturalistic reading is more challenging than modelling passive speech listening. Unlike passive listening, naturalistic reading involves internally-paced eye movements that determine when and for how long visual information is acquired. This results in variable sampling rates that make it difficult to temporally align continuous stimulus representations with ongoing neural activity in real time. Consequently, many previous EEG/MEG studies of reading have relied on either time-paced rapid serial visual presentation, in which words are displayed rapidly and sequentially at fixed intervals (e.g., Frank et al., 2015; Soghoyan et al., 2025), or self-paced reading paradigms, which simplify time-locking to each word through button presses (e.g., Ditman et al., 2007). While these approaches facilitate neural alignment to eye fixations, they constrain natural eye-movement patterns (e.g., variable fixation rates, regression patterns) and limit ecological validity (Dimigen et al., 2011; Radach et al., 2008; Rayner et al., 1989). Moreover, unlike speech, text does not unfold as a continuous waveform, but is sampled through discrete fixation inputs disconnected by saccades (Rayner, 1998). Here, we address these methodological challenges by time-locking linguistic predictors to fixation onsets and durations, we transform discrete sampling events into a time series representation approximating continuous input (i.e., pseudo-continuous regressors) compatible with mTRF analysis. This allows neural tracking of linguistic features to be examined under more naturalistic reading conditions.

Using the above-described approach of modelling brain responses during naturalistic reading, we addressed our main question through two sub-aims. First, we examined whether, and to what extent, neural tracking of different features was modulated across task goals. Second, we tested whether neural tracking of these linguistic features was associated with changes in behavioural performance (i.e., the extent to which participants achieved the assigned goal) under specific goal conditions. We focused on three levels of linguistic processing at the orthographic, lexical, and contextual-semantic levels:

1. Neural tracking of orthographic information was expected to be enhanced for the Proofreading goal, reflecting increased sensitivity to lower-level word-form detail. In contrast, we expected orthographic tracking to remain stable or potentially reduced in the Comprehension goal condition compared to Baseline, as orthographic information would be less directly relevant to the task goal.
2. Word Frequency tracking can be expected to increase due to deeper lexical processing when attending to semantic details in the Comprehension condition (e.g., Radach et al., 2008), or to remain stable or potentially decrease if reliance on higher-level contextual information potentially modulates reliance on lexical processing (Levy, 2008; Norris, 2006).
3. We predicted that the Comprehension goal would lead readers to prioritise higher-level semantic and discourse coherence, thereby enhancing neural tracking of contextual-semantic information in this condition compared to Baseline. On the other hand, global coherence would be less task-relevant during the Proofreading condition, and therefore contextual-semantic tracking was expected to remain the same or be reduced compared to Baseline.

## Method

### Participants

Forty participants (26 female, 14 male; *M*_Age_ = 19.7 years, *SD*_Age_ = 2.0, range: 18-26) with normal vision and no language impairments were recruited through the University of Aberdeen’s SONA participant pool and word-of-mouth. All were native speakers of English (learned before the age of 5), and 20 reported speaking at least one additional language proficiently. This study received ethical approval from the Psychology Ethics Committee at the University of Aberdeen.

### Materials

Three Indonesian legends in English language were adapted from web (https://indonesianfolklore.blogspot.com/2007/12/timun-mas-folklore-from-central-java.html) and expanded using ChatGPT-4o (OpenAI, 2025), then edited to equate story length. Each story was divided into 16 passages, with a mean of 104.8 words per passage (*SD* = 13.3, range: 75-131). For each story, two versions were created. The first version contained no spelling errors and was used in the Baseline and Comprehension goal conditions. The second version was created by randomly introducing three misspelled words amongst content words within each passage for the Proofreading goal condition (e.g., mirrors, mirorrs; crimson, crimsen). For each passage, two three-option multiple-choice questions were generated: one proofreading question and one comprehension question, for the Proofreading and Comprehension conditions, respectively. Proofreading questions asked: “Which of these words were misspelled in the section of the story that you just read?”, whereas Comprehension questions asked specific details from the preceding passage, such as: “What was the Harp made of?”. In total, this resulted in three stories, each with 16 passages, two text versions, and corresponding Proofreading and Comprehension questions.

Stimulus features were extracted to represent linguistic features at the orthographic, lexical, and contextual-semantic levels:

1. Orthographic neighbourhood size (ONS; Coltheart et al., 1977), calculated as the number of words differing from the target word by one letter (e.g., cat and can), was included as an index of lower-level, predominantly sub-lexical, processing.
2. Word frequency (Zipf values, ranging from 1-7 on a log10 scale), were obtained from the SUBTLEX-UK lexicon (van Heuven et al., 2014), was included as an index of lexical processing, reflecting the base probability of encountering the target word regardless of the context in which it appears.
3. Word Surprisal, calculated as the negative log probability of the target word given its preceding context, was estimated using GPT-NeoX-2B (Black et al., 2022) and included as an index of higher-level contextual-semantic processing. This quantifies how unexpected a word is given the preceding context, with lower values indicating higher contextual predictability (Levy, 2008). Although typically correlated with lexical frequency (e.g., Shain, 2019), word surprisal additionally captures the extent to which the word is (un)expected within the discourse context, with additive and dissociable effects compared to frequency (e.g., Goodkind & Bicknell, 2021; Huizeling et al., 2022; Shain, 2024; Staub & Benatar, 2013). See Appendix A for correlations between predictors.

### Procedure

Participants completed three blocks, each corresponding to one condition. In a within-subjects design, participants always first completed the Baseline goal condition, followed by Proofreading and Comprehension goal conditions in a counterbalanced order. This ordering ensured that goal manipulation did not influence baseline reading in the first block. The three story narratives were assigned to the three conditions in a counterbalanced order across participants (i.e., 2×3: six total pairings).

Before each block, participants were instructed to read silently at their own pace and while minimising head movements. In the Proofreading condition, they were asked to proofread the text and told that they would later be tested on spelling mistakes. In the Comprehension condition, participants were instructed to pay attention to the story content and told that they would later be tested on specific facts or events. They were not given further instructions for the Baseline condition and were simply instructed to read as they normally would.

Participants were seated 79 cm away from the display monitor (Display++ 32” LCD, Cambridge Research Systems; 1920 × 1080p, 120 Hz, 71.0 x 39.5 cm) with their head stabilised using a head rest. Neural activity was recorded using 64-channel (Ag/AgCl electrodes) BioSemi ActiveTwo EEG system sampled at 512 Hz. Eye movements were recorded from the left eye using the EyeLink 1000+ eye-tracking system (SR Research Ltd., 2013) at a sampling rate of 1000 Hz. The eye-tracker was calibrated using the default EyeLink 9-point calibration procedure, with validation errors kept below 1° of visual angle. Drift correction at the top of screen was performed before each passage, and re-calibration was performed before each block.

Each narrative was presented as a series of 16 passages. Passage screens were presented in pairs, followed by two task-related questions, after which the next pair of passages was shown (e.g., passage 1, passage 2, question 1, question 2, …). Each passage started with a gaze-contingent black fixation dot (20 pixels) located at the top left corner of the screen at [300, 125] on a white screen. Participants were required to fixate within 45 pixels around the centre of the dot for at least 200 ms for the passage to appear. Passages were presented in black Times New Roman font (size 14, triple line spacing) on a white background. Participants pressed the space bar to proceed to the next screen. Between passages, a blank interval screen containing “…” at the centre was presented for 3000 ms. Task-related questions were also presented in pairs in the form of multiple-choice questions. Participants responded by pressing [1], [2], or [3] on the keyboard. In the Baseline condition, no questions were presented, but instead participants were prompted to press [1] to continue.

### Analysis

#### Eye movement data preprocessing

Eye movement data were examined manually offline using the EyeLink Data Viewer software (Version 4.4.1; SR Research Ltd., 2024). Fixation positions were corrected by uniformly shifting all fixation coordinates together (without moving individual points) if misalignments or drifts were detected. These were then processed using a custom script in MATLAB (MathWorks, 2022) to calculate per-passage (display screen) reading times and regression rates, and per-word fixation times. Reading times were calculated as the total length of time between the start of display and the button press indicating that the participant finished reading. Regression rates were calculated by dividing the number of regressions by the total number of fixations. Mean per-word fixation times were calculated across all fixated words.

#### EEG preprocessing

EEG data were preprocessed using MNE-Python (Gramfort et al., 2013). Data were first manually inspected to identify bad channels, then average re-referenced. Independent Component Analysis (ICA) was performed (using good channels only) to remove non-neural artifacts such as blinks. Previously marked bad channels were then interpolated using data from neighbouring electrodes. Finally, the signal was filtered using a zero-phase Finite Impulse Response (FIR) band-pass filter of 0.2-40 Hz.

For each participant, EEG data were segmented from the onset of each passage display to each space-bar response (marking the end of that passage). These were z-score normalised, then concatenated for each condition, with 200 ms zero-padding between segments.

#### mTRF analysis

Subsequent analyses were conducted in MATLAB (MathWorks, 2022).

Eye-tracking data were used to align neural activity with linguistic features of the currently fixated word. Stimulus features were modelled as a boxcar function (i.e., each value spans the whole duration of fixation) to form pseudo-continuous regressors that reflected the temporal progression of input. These were z-score normalised, then down-sampled (decimated) to 128 Hz alongside the EEG data.

Multivariate Temporal Response Function (mTRF) modelling was conducted using the mTRF toolbox (Crosse et al., 2016; Crosse et al., 2021). Forward modelling (i.e., predicting neural data from stimulus features) was used to model the neural tracking of multivariate stimulus features: ONS, Zipf, and Surprisal. Fixation onsets were included as a control regressor to reduce eye movements confounds (i.e., for each unique fixation). Models were run separately for each participant and Condition, with a temporal lag window of [-100, 600] ms. The optimal lambda (regularisation parameter) was estimated across the logarithmic range from 10^-7^ to 10^7^ through a 10-fold cross-validation procedure (mTRFcrossval). The final model was then fitted using the obtained lambda through a 10-fold training and testing procedure (mTRFtrain, mTRFpredict).

mTRF calculates the prediction accuracy (R-values) between predicted and the observed EEG data, which we used as an index estimating the strength of neural encoding of individual features. To estimate the contribution (and the tracking strength) of specific linguistic features to the overall model fit, for each stimulus feature of interest (ONS, Zipf, Surprisal) we use the partial R method (Crosse et al., 2021). For each feature of interest corresponding regressor was circularly shifted 100 times to generate a null distribution that preserved its temporal structure while disrupting the phase relationship between the feature and neural signal (Bialek & de Ruyter van Steveninck, 2005; Crosse et al., 2021). The shifted feature vector was then added to other unaltered features to derive permuted model fit R-values. The partial R-value, indexing feature-specific neural encoding, was then obtained by subtracting the full model R-values by the specific-feature permuted R-values. Higher partial R-values were therefore interpreted as stronger neural tracking of that feature after accounting for the other predictors included in the model. TRF model weights are reported in Appendix B.

The use of mTRF helps address the problem of overlapping neural responses during naturalistic reading, that is, both between consecutive fixations, and between neural responses to reading events and neural artefacts not associated with linguistic feature processing (see Dimigen & Ehinger, 2021). As naturalistic eye movements occur rapidly and irregularly, neural responses to one word may still be unfolding when the next word is fixated. However, the time-lag window used in mTRF allows delayed responses to each word fixation to be estimated over time, which would reduce the extent to which neural responses from overlapping neighbouring fixations are misattributed to the currently fixated word. In addition, including fixation onsets as a control regressor into mTRF allows fixation-related activity to be modelled separately. This is important as eye movements themselves generate neural responses and artefacts that may overlap with linguistic processing (Dimigen & Ehinger, 2021; Ehinger & Dimigen, 2019; Plöchl et al., 2012). While this does not fully eliminate eye-movement artefacts or isolate linguistic processing perfectly, mTRF provides a useful deconvolution approach for separating the unwanted contribution of EEG signals not directly related to our research question by accounting for fixation-related activity during naturalistic reading.

### Analysis approach

All statistical analyses were conducted in RStudio (Version 2023.12.1.402; Posit Team, 2024). Linear and generalised linear mixed-effects models were fitted using the lme4 and lmerTest packages (Bates et al., 2015; Kuznetsova et al., 2017).

Analysis of eye movement data was conducted for each metric (reading time, regression rate, word fixation) using linear mixed-effects models. Condition was dummy coded with Baseline as the reference level for direct comparisons between Proofreading vs. Baseline and Comprehension vs. Baseline. The full model was Eye_movement ∼ Condition + (1 | Participant).

For each stimulus feature, partial R-values were averaged across all EEG channels per participant and Condition. These mean R-values were used to index the strength of neural tracking for ONS, Zipf, and Surprisal separately. Two sets of analyses were conducted to address the study aims.

We first examined whether neural tracking of each feature differed across goal conditions. Feature-specific partial R-values were modelled using linear mixed-effects modelling, with Condition as a fixed effect. Separate models were fitted for ONS, Zipf, and Surprisal tracking. Similar to our eye movement analysis, Condition was dummy coded with Baseline as the reference level for direct comparisons between Proofreading vs. Baseline and Comprehension vs. Baseline. The full model was Feature_Rval ∼ Condition + (1 | Participant).

Next, we examined whether neural tracking of specific linguistic features predicted behavioural performance (i.e., whether they achieved the task goal) in the two goal conditions. As behavioural questions were only given in the Proofreading and Comprehension conditions, the Baseline condition was excluded from this analysis. Accuracy was modelled using generalised linear mixed-effects modelling as a function of all the partial R-values (z-scored). The full model was Condition_Accuracy ∼ Rval_surprisal_z + Rval_zipf_z + Rval_ONS_z + (1 | Participant) + (1 | Question). No random slopes were included due to convergence problems.

## Results

### Eye movements

Examination of the eye movement data (Table 1) showed expected differences (e.g., Daneman et al., 1995; Kaakinen & Hyönä, 2010; Schotter et al., 2014) in reading times, regression rates, and word fixations across conditions: on average, per-passage reading times and per-word fixation times were significantly longer in the Proofreading condition compared to the Baseline no-goal condition, but not between Comprehension and Baseline conditions.

**Table 1.**
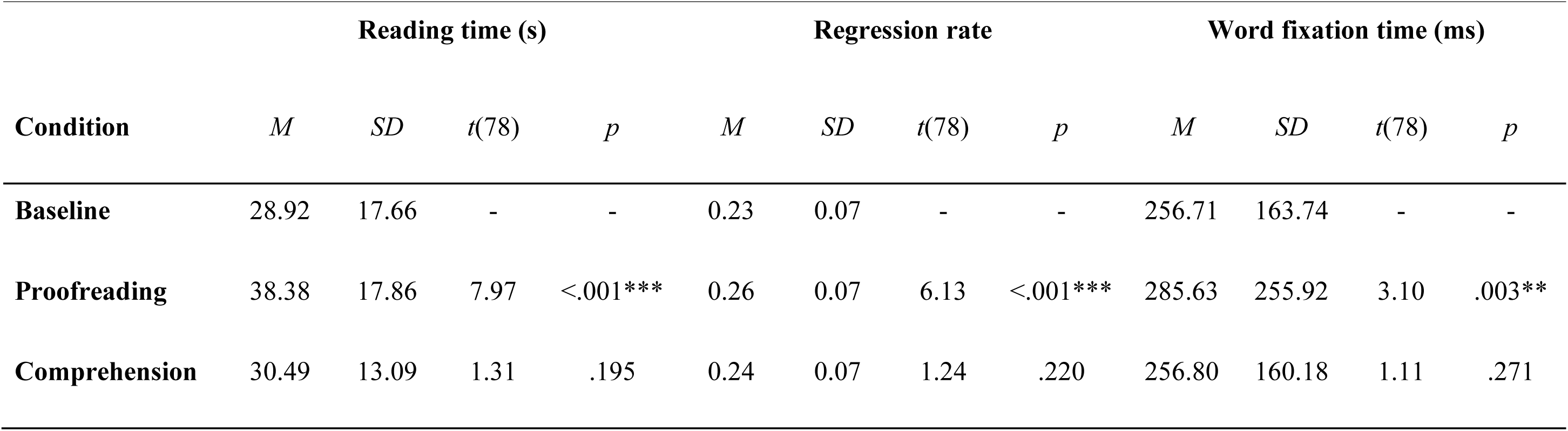
Analysis of eye movement data. Reading times and regression rates were calculated per display screen. Mean per-word fixation times were calculated across all fixated words. Linear mixed-effects models were run on each measure: *t* and *p*-values show the planned contrasts for the Proofreading vs. Baseline and Comprehension vs. Baseline conditions.

As our primary question relates to goal-related modulation in neural tracking of linguistic features, we focus subsequent analyses on the eye-movement aligned neural data.

### Neural tracking

Figure 1 shows average participant-level partial R-values along with the mean scalp topographies for ONS, Zipf, and Surprisal tracking across conditions. Decoding in all conditions was significantly above chance at the group level (see Appendix C).

**Figure 1.**
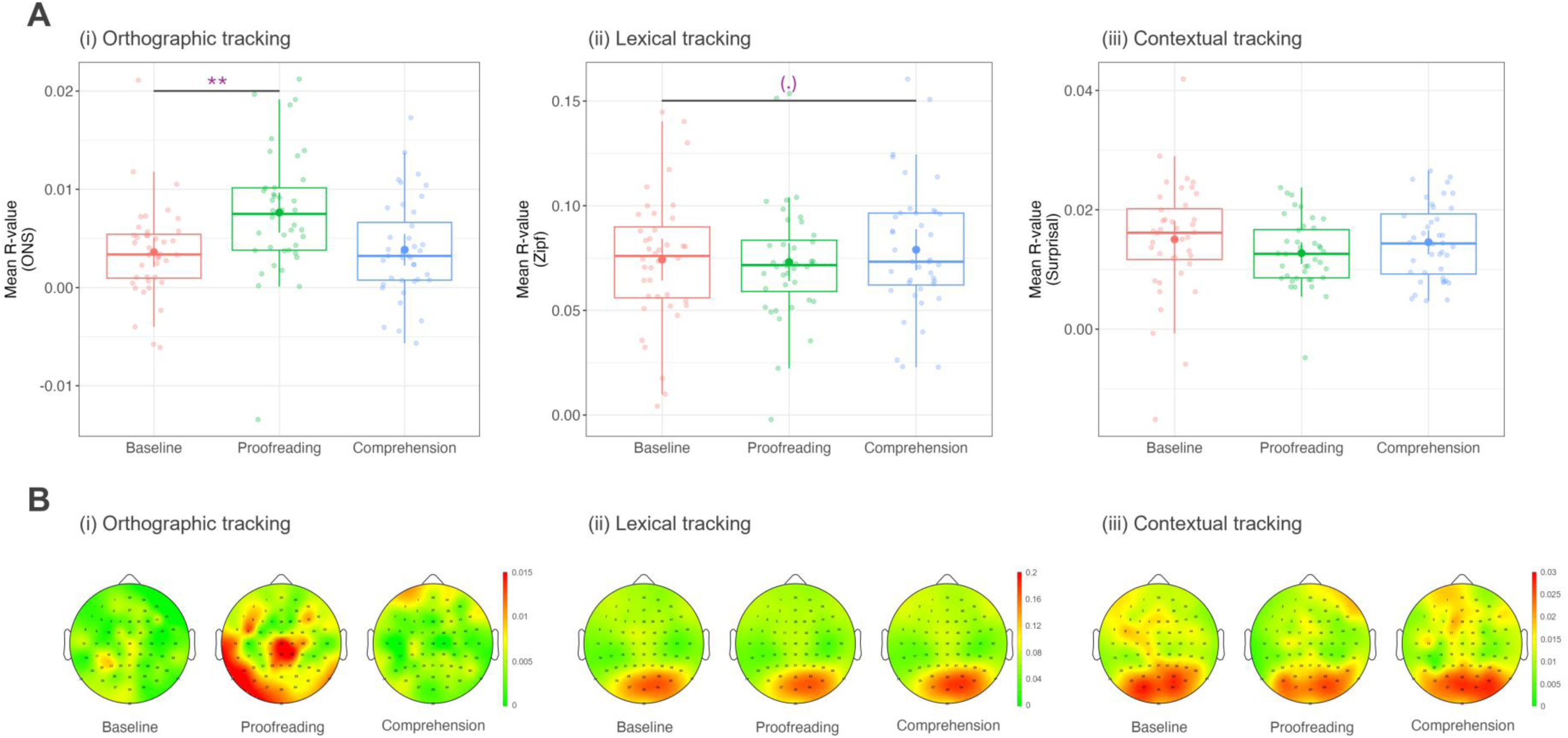
Partial R-values across conditions. (A) Distribution of participant prediction correlation means (Partial R-values) of predictors (i) Orthographic (ONS), (ii) Lexical (Zipf Frequency), and (iii) Contextual (Semantic Surprisal), averaged across all electrodes. (B) Mean scalp topographies averaged across participants.

To address our first aim, we examined the effect of Condition on Feature tracking by fitting three separate linear mixed-effects models with ONS, Zipf, and Surprisal partial R-values as the dependent variables respectively. Model intercepts were significant for all three features: ONS, *β* = 0.004, *SE* = 0.001, *t*(117.0) = 4.20, *p* < .001; Zipf, *β* = 0.074, *SE* = 0.005, *t*(46.9) = 15.59, *p* < .001; Surprisal, *β* = 0.015, *SE* = 0.001, *t*(105.6) = 12.83, *p* < .001. This demonstrates that all linguistic properties being examined were reliably tracked (significantly above zero) in the Baseline condition. Comparing conditions, there were three findings of note: First, ONS tracking was significantly higher in the Proofreading than in the Baseline condition, *β* = 0.004, *SE* = 0.001, *t*(117.0) = 3.32, *p* = .001, but did not differ significantly between the Comprehension and Baseline conditions, *β* = 0.0002, *SE* = 0.0012, *t*(117.0) = 0.20, *p* = .842. This suggests that neural tracking of orthographic information was selectively increased only when relevant (i.e., when participants had a proofreading goal). Next, Zipf frequency tracking was marginally higher in the Comprehension compared to the Baseline condition, *β* = 0.005, *SE* = 0.002, *t*(78.0) = 1.90, *p* = .061, but was comparable between the Proofreading and Baseline conditions, *β* = −0.001, *SE* = 0.002, *t*(78.0) = −0.47, *p* = .640. This trend is consistent with increased neural tracking of lexical information in the Comprehension goal. Finally, Surprisal tracking did not differ significantly between the Proofreading and Baseline conditions, *β* = −0.002, *SE* = 0.001, *t*(78.0) = −1.60, *p* = .115, nor between the Comprehension and Baseline conditions, *β* = −0.0005, *SE* = 0.0014, *t*(78.0) = −0.33, *p* = .744, providing no evidence that goals significantly modulated neural tracking of contextual-semantic processing. See Appendix D for additional pairwise post-hoc comparisons between conditions.

### Correlation between neural tracking and behavioural scores

Average behavioural accuracy was 72.2% (*SD* = 15.2) in the Proofreading condition (detecting spelling errors) and 85.5% (*SD* = 10.9) in the Comprehension condition (answering comprehension questions).

To address our second aim, we examined whether feature-specific neural tracking during reading predicted later behavioural performance within each goal condition (Proofreading and Comprehension) using generalised linear mixed-effects modelling (Condition_Accuracy ∼ Rval_surprisal_z + Rval_zipf_z + Rval_ONS_z + (1 | Participant) + (1 | Question)). In the Proofreading condition, only ONS tracking significantly predicted accuracy, *β* = 0.243, *SE* = 0.121, *z* = 2.00, *p* = .045 (see Figure 2), suggesting that stronger orthographic tracking contributed to better performance when participants aimed to detect spelling errors. Zipf and Surprisal tracking did not contribute significantly to Proofreading accuracy, *β* = 0.102, *SE* = 0.135, *z* = 0.75, *p* = .451, and *β* = −0.175, *SE* = 0.136, *z* = −1.29, *p* = .197, respectively. On the other hand, in the Comprehension condition, Zipf tracking significantly predicted accuracy, *β* = 0.331, *SE* = 0.161, *z* = 2.06, *p* = .039 (see Figure 2), while Surprisal tracking showed a marginal effect, *β* = 0.286, *SE* = 0.156, *z* = 1.83, *p* = .068. This suggests that stronger lexical tracking, and perhaps contextual-semantic tracking (weakly), contributed to greater attention to specific details of facts or events whilst reading. ONS tracking did not significantly predict accuracy in the Comprehension condition, *β* = −0.143, *SE* = 0.147, *z* = −0.97, *p* = .330.

**Figure 2.**
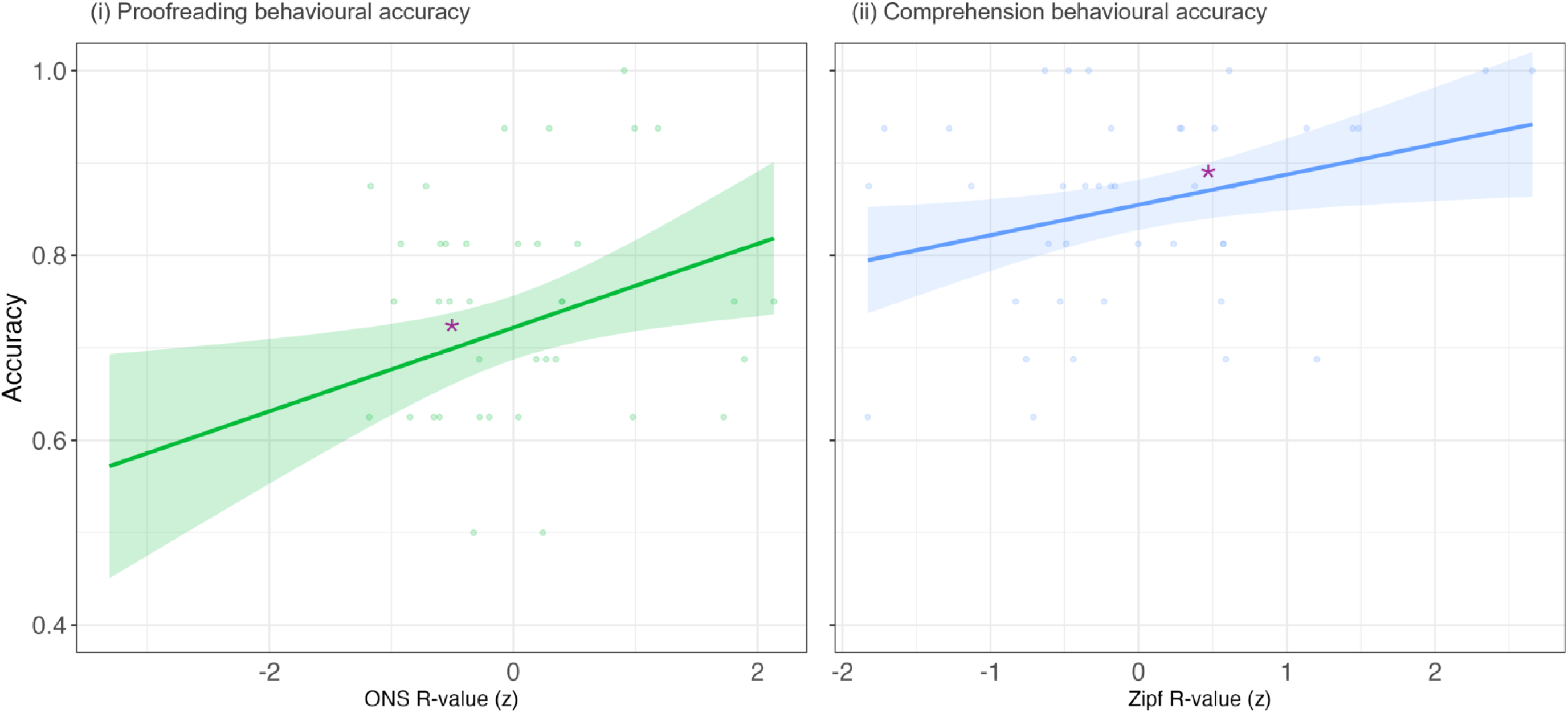
Behavioural scores in Proofreading and Comprehension conditions as predicted by neural tracking of linguistic properties Orthographic Neighbourhood Size (ONS) and Lexical Frequency (Zipf).

## Discussion

By combining EEG with eye-tracking, the present study examined how short-term task goals modulate neural tracking of linguistic features during fully naturalistic reading. Our findings demonstrate that task goals are able to selectively alter the weighting of linguistic information analysis in a goal-relevant manner. (1) At the lower orthographic level, neural tracking of ONS was significantly enhanced during Proofreading but remained stable during Comprehension relative to Baseline. (2) At the lexical level, neural tracking of Zipf word frequency showed weak evidence (non-significant trend) of enhancement during Comprehension but not during Proofreading relative to Baseline. (3) Finally, at the higher contextual-semantic level, neural tracking of Surprisal was not significantly modulated by task goals. This pattern aligns with participants’ behavioural performance: ONS tracking significantly predicted Proofreading accuracy, whereas Zipf tracking significantly predicted Comprehension accuracy. These findings suggest that rather than simply increasing attention or effort globally, goals modulate the relative weighting of linguistic levels in order to optimise linguistic processing at the levels that contribute to achieving the specific goal.

Goal-dependent modulation was most evident at the orthographic level. When participants were instructed to detect spelling errors, neural tracking of ONS increased relative to Baseline reading. This suggests that Proofreading enhanced sensitivity to lower-level word-form detail, consistent with the idea that spelling error detection relies on monitoring of word-form detail for letter-by-letter orthographic comparison between the perceived form and expected lexical representation (Norris, 1984; O’Connor & Forster, 1981). This finding is also supported by the behavioural analysis, which showed that participants who tracked orthographic information more strongly were better able to detect spelling errors, suggesting that enhanced orthographic tracking contributed directly to successful goal performance. In contrast, orthographic tracking did not appear to be modulated during the Comprehension condition relative to Baseline. This may be because orthographic processing remains necessary for visual word recognition and subsequent higher-level processes, and may not need to be actively or explicitly reduced to support more focused contextual analysis. This extends previous behavioural work showing that proofreading tends to produce slower and more careful reading with increased tracking of/sensitivity to word-form information, as indicated by increased word length effects (Kaakinen & Hyönä, 2010), and, as in our own data, longer reading times.

At the lexical level, lexical frequency tracking showed a weaker pattern of modulation across conditions. Lexical frequency showed the trend of increasing tracking (i.e., marginally higher) in the Comprehension than the Baseline condition; however, lexical tracking did significantly predict Comprehension goal accuracy. This pattern of results suggest that lexical-level processing contributed to successful performance in this condition. It may be that deeper semantic processing in general occurs when participants were explicitly instructed to read for comprehension (e.g., Radach et al., 2008). However, if this were the case, we arguably should have observed contextual-semantic tracking to also increase alongside frequency tracking. Alternatively, although we initially predicted that Comprehension goal would primarily enhance contextual-semantic tracking, one possibility is that readers focused more on lexical detail than global contexts in anticipation of the comprehension questions, which may explain why lexical tracking predicted successful goal achievement more strongly than contextual-semantic tracking. However, this interpretation should be taken cautiously, as participants did not know the specific questions in advance.

On the other hand, lexical frequency tracking was not significantly modulated during the Proofreading condition, nor did it predict Proofreading accuracy. This is contrary to previous studies reporting increased frequency effects during proofreading, which instead suggest modulation at the lexical level when detecting errors (Daneman et al., 1995; Kaakinen & Hyönä, 2010; Schotter et al., 2014). This was unexpected, as error detection is expected to involve accessing lexical representations in addition to orthographic comparisons (Foster, 1976; Norris, 1984; O’Connor & Forster, 1981), thus supporting error detection alongside orthographic tracking (e.g., Kaakinen & Hyönä, 2010; Perea et al., 2005; Schotter et al., 2014). However, the behavioural analysis found that successful Proofreading performance relied more strongly on orthographic than lexical tracking. This finding, together with the lack of observable modulation at the lexical level during Proofreading, suggests that the level of lexical processing normally engaged during standard reading is generally sufficient in supporting lexical access to the expected word form for successful error detection, with the limiting factor being orthographic monitoring.

Similarly, the absence of a significant goal-dependent modulation at the contextual-semantic level was unexpected based on our prediction that Comprehension would enhance contextual-semantic processing. Firstly, this may be because Baseline reading without explicit goal instructions may already have involved strong contextual-semantic processing, as this is the dominant ecological function of reading. The adult reading system is likely optimised through lifelong experience for extracting semantic and discourse-level information under default conditions (e.g., Deniz et al., 2019; Swinney, 1979), and thus participants would have naturally read for meaning even without explicit instructions (i.e., defaulting to comprehension goal under baseline conditions). Thus, top-down contextual processing may persist across task goals. In this case, there would be limited room for Surprisal tracking to increase further when explicitly instructed to attend to content. An explicit comprehension goal would thus reflect similar processing patterns as in typical no-goal reading states, whereas a proofreading goal would require a more pronounced re-allocation of precision weightings towards lower-level orthographic features.

Additionally, semantic and discourse processing may be less readily modulated by short-term goals as predicted by the predictive processing framework. Higher-level representations are shaped by longer-term reading experiences and are thus more resilient to more transient fluctuations in the environment (Kiebel et al., 2008). Given the transient nature of task goals, adaptation is then expected to occur rapidly while maintaining stability of higher-order representations. Lower-level representations (e.g., visual, orthographic) would hence be more readily re-weighted under transient demands, whereas higher-level processing (e.g., semantic, discourse) would remain relatively stable as they would be shaped by longer-term experiences. This account also explains the pattern of our findings, in which the orthographic level was most readily modifiable (being the lowest level in the present study), then lexical processing being more weakly modulated (mid-level), and contextual-semantic processing as the most stable (highest level).

Surprisal tracking was also not modulated or specifically reduced in the Proofreading condition. This suggests that participants continued to process and track global coherence even when only attempting to detect spelling errors, maintaining higher-level linguistic processing while additionally enhancing orthographic tracking. This is consistent with evidence that readers do not merely scan text for spelling errors while proofreading, but also continue to monitor semantic processing (e.g., Levy & Begin, 1984). Proofreading may therefore involve enhanced processing of orthographic information on top of normal reading processes rather than replacing or suppressing higher-level linguistic processing. More broadly, none of the selected linguistic features showed clear evidence of goal-dependent down-regulation. This is important as it may suggest that modulation operated primarily through selective enhancement of goal-relevant information rather than an active suppression for information that was less directly relevant. One speculative interpretation could be that reading has become an optimised and flexible process such that baseline processing is already operating at optimal levels without utilising all available processing resources. However, this needs to be carefully interpreted and should not be taken as evidence that processing resources are unlimited or that no down-weighting occurred globally. The selected predictors may simply have been more sensitive to enhancement than suppression, or have captured processes that remain necessary across all goal conditions.

While research on adaptive neural mechanisms in reading is sparse, there is substantial evidence for adaptive analysis of linguistic features in speech. Firstly, selective attention paradigms have shown that neural representations of attended speech are preferentially encoded compared to unattended input at both low-level acoustic (e.g., speech envelope) and higher-level linguistic (e.g., lexical, semantic) stages (e.g., Broderick et al., 2018; Jaeger et al., 2025; Mesgarani & Chang, 2012; Zion Golumbic et al., 2013; Olguin et al., 2018; O’Sullivan et al., 2015), demonstrating robust effects of top-down attentional modulation. Next, when uncertainty is selectively manipulated at a particular processing level with other aspects of input held constant, neural tracking does not change uniformly across the hierarchy (e.g., Karunathilake et al., 2023; Piazza et al., 2025). Instead, modulation appears to be targeted to the source of uncertainty, with stronger disruptions in acoustics generating cascading effects for higher levels. For instance, Karunathilake et al. (2023) utilised a vocoded speech paradigm to manipulate the intelligibility of speech while preserving its acoustic features across vocoded items, and found that word-level tracking varied with comprehension whereas low-level acoustic tracking remained relatively unaffected. Conversely, when acoustic uncertainty is introduced (e.g., through speaker variability or introducing temporal discontinuity) while semantic content is preserved, lower-level phonemic representations are selectively enhanced (Klimovich-Gray et al., 2021; Piazza et al., 2025). Tracking of contextual semantic information is sensitive to the degree of disruption of lower-level acoustic information, with more subtle disruptions (speaker variability) showing preserved tracking (Piazza et al., 2025) and stronger disruptions (temporal discontinuity) weakening higher-order analysis (Klimovich-Gray et al., 2021). Such dissociation suggests that specific linguistic levels can be selectively up- or down-weighted depending on the degree and location of the uncertainty within the speech stream. Therefore, our findings in goal-oriented reading add to the language processing literature more broadly, suggesting that flexible weighting of linguistic features across the processing hierarchy may be a unified mechanism across spoken and written language modalities that operates in response to both external stimulus-driven demands and internally generated goals.

## Conclusion

The present study provides first evidence that task goals selectively modulate neural tracking of lower-level orthographic features during naturalistic reading, consistent with the precision-weighted account of predictive language processing. Critically, goals did not produce a general increase in neural tracking across all linguistic features, nor did they clearly suppress goal-irrelevant information. Instead, we saw that targeted modulation occurred for the feature most directly relevant to the proofreading task. When detecting spelling errors, orthographic tracking was selectively enhanced, which consequently predicted error detection accuracy. In contrast, comprehension accuracy was predicted by tracking local lexical information, while top-down contextual-semantic tracking was maintained across all reading conditions. These findings suggest that goal-oriented reading is characterised by the selective re-weighting of linguistic information at the lexical and sub-lexical level towards those that contribute the most to successful goal attainment. By combining eye-tracking with EEG, we have thus shown that goals modulate the neural tracking of specific linguistic features online during naturalistic reading. Future work combining eye-tracking and mTRF modelling should explore further linguistic features that are beyond the scope of the current study, such as entropy-based measures (Shannon, 1948) like orthographic uncertainty (Westbury & Yang, 2024) and semantic entropy (Kuhn et al., 2023), and linguistic structure such as part-of-speech syntactic probability. Moreover, in addition to fixation onsets, future work could include additional oculomotor covariates (e.g., saccade amplitudes, incoming direction, pupil diameter) using non-linear deconvolution approaches, such as spline regression (Dimigen & Ehinger, 2021). Collectively, these promising new approaches will allow us to make progress towards a more explicit neurocognitive account of how processing levels are dynamically modulated during naturalistic reading.

## Acknowledgements

This research was supported by the Economic and Social Research Council (ESRC) and the Scottish Funding Council (SFC) via the Scottish Graduate School of Social Science (SGSSS) Advanced Quantitative Methods (AQM) doctoral training mechanism (PA, BT, AK, JS).

AKG acknowledges support and networking opportunities provided by Scottish Imaging Network: A Platform for Scientific Excellence (SINAPSE). JS acknowledges funding from the Juan de la Cierva Postdoctoral Fellowship programme 2025-2026, the Basque Government through the BERC 2022-2025 programme, and the Spanish State Research Agency through BCBL Severo Ochoa excellence accreditation CEX2020-001010/AEI/10.13039/501100011033. We thank Marta Brzeska and Catherine Mulder for help with EEG data collection.

## Author contributions

Pepita Alex: Conceptualization, Investigation, Data curation, Formal analysis, Methodology, Visualization, Writing – original draft

Julia Schwarz: Methodology, Supervision, Writing – review and editing

Ben Tatler: Conceptualization, Funding acquisition, Methodology, Supervision, Writing – review and editing

Agnieszka Konopka: Conceptualization, Funding acquisition, Methodology, Supervision, Writing – review and editing

Anastasia Klimovich-Gray: Conceptualization, Data curation, Funding acquisition, Methodology, Project administration, Supervision, Writing – review and editing

## Appendix A Pearson Correlations Between Linguistic Predictors

| Measure | 1 | 2 | 3 |
| --- | --- | --- | --- |
| 1. ONS | - | - | - |
| 2. Zipf | .49 | - | - |
| 3. Surprisal | -.19 | -.36 | - |

## Appendix B: TRF Model Weights in the Occipital-Parietal Region

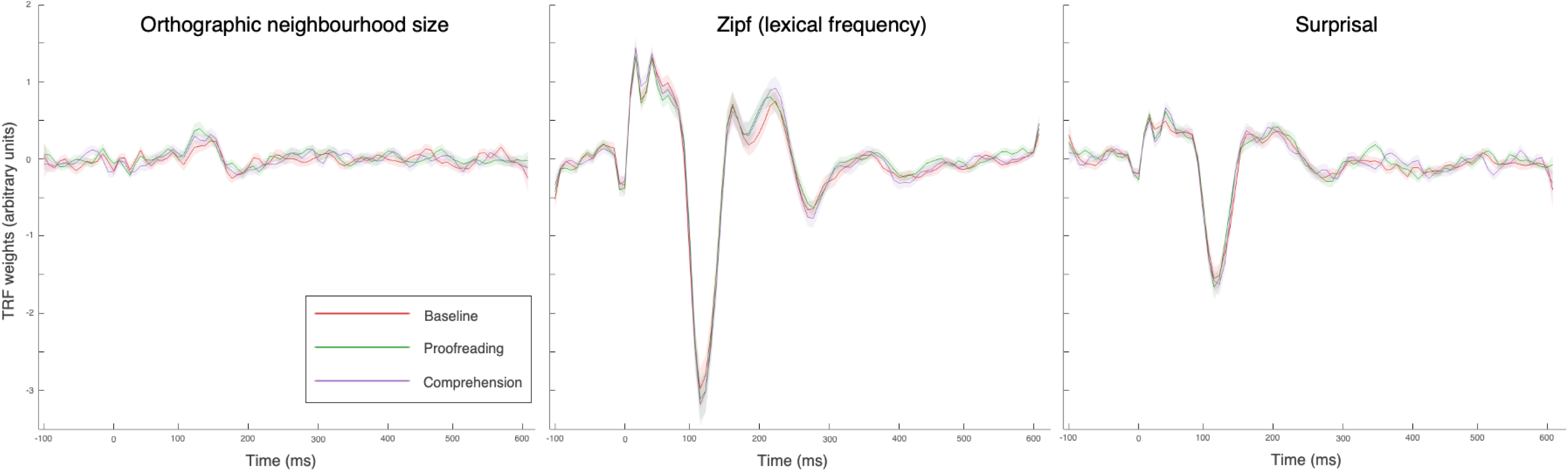

## Appendix C: T-tests of Partial R-values Against Zero

| **Condition** | **ONS** |  | **Zipf** |  | **Surprisal** |  |
| --- | --- | --- | --- | --- | --- | --- |
|  | *t*(39) | *p* | *t*(39) | *p* | *t*(39) | *p* |
| **Baseline** | 4.8 | < .001*** | 15.01 | < .001*** | 9.83 | < .001*** |
| **Proofreading** | 7.58 | < .001*** | 16.34 | < .001*** | 14.27 | < .001*** |
| **Comprehension** | 4.84 | < .001*** | 16.27 | < .001*** | 14.64 | < .001*** |

*Note.* One-sided, one-sampled t-tests against zero.

## Appendix D: Pairwise Post-hoc Comparisons Between Conditions

**Appendix D** **Pairwise Post-hoc Comparisons Between Conditions**
| **Contrast** | **ONS** |  | **Zipf** |  | **Surprisal** |  |
| --- | --- | --- | --- | --- | --- | --- |
|  | *t*(78) | *p* | *t*(78) | *p* | *t*(78) | *p* |
| **Baseline - Proofreading** | -3.32 | .004** | 0.47 | .640 | 1.60 | .031 |
| **Baseline - Comprehension** | -0.20 | .842 | -1.90 | .092 | 0.33 | .744 |
| **Proofreading - Comprehension** | 3.12 | .004** | -2.37 | .061 | -1.27 | .031 |

*Note.* Degrees of freedom were calculated using the Kenward-Roger method, and *p*-values were adjusted for multiple comparisons using the false discovery rate (FDR) method across three tests.

